# Integrity assay for messenger RNA in mouse and human brain samples and synaptosomal preparations

**DOI:** 10.1101/2024.01.12.575353

**Authors:** Daina Bujanauskiene, Kajus Merkevicius, Ugne Kuliesiute, Jaroslav Denkovskij, Simonas Kutanovas, Gediminas Luksys, Saulius Rocka, Eiva Bernotiene, Urtė Neniskyte

**Affiliations:** VU LSC-EMBL Partnership for Genome Editing Technologies, Life Sciences Center, Vilnius University, Vilnius, Lithuania; Institute of Biosciences, Life Sciences Center, Vilnius University, Lithuania; Clinic of Paediatrics, Institute of Clinical Medicine, Faculty of Medicine, Vilnius University, Vilnius, Lithuania; Department of Regenerative Medicine, Center for Innovative Medicine, Vilnius, Lithuania; Center of Neurosurgery, Vilnius University Hospital Santaros Klinikos, Vilnius, Lithuania; Clinic of Neurology and Neurosurgery, Faculty of Medicine, Vilnius University, Vilnius, Lithuania

**Keywords:** RNA integrity, messenger RNA, ribosomal RNA, 5’:3’ assay, RT-qPCR, RIN, synaptosomes

## Abstract

Traditionally, RNA integrity evaluation is based on ribosomal RNAs (rRNAs). Nevertheless, gene expression studies are usually focused on protein coding messenger RNAs (mRNAs). As rRNA and mRNA have significant structural and functional differences, the assumption that rRNA integrity properly represents mRNA integrity may not be accurate. Moreover, contrary to whole tissue RNA samples, subcellular preparations such as synaptosomes contain almost no rRNA, thus prohibiting the use of traditional rRNA-based methods to assess sample RNA integrity. Here we present a RT-qPCR based assay, which estimates mRNA integrity by comparing the abundance of 3’ and 5’ mRNA fragments in a long constitutively expressed mouse or human *PGK1* mRNA. The assay was tested and validated using plasmids with cloned 3’- and 5’-ends of the *PGK1* cDNA reflecting different ratios of 3’ and 5’ cDNA amplicons in partially degraded RNA samples. The accuracy of integrity score calculation was ensured by integrating a mathematical correction of qPCR results to account for the variable amplification efficiency of different primer pairs. The 5’:3’ assay was used to quantify RNA degradation in heat-degraded mouse and human brain tissue RNA as well as in clinical human brain RNA samples. Importantly, the expression of housekeeping genes correlated better with 5’:3’ integrity value than with the RIN. Finally, we were even able to use 5′:3′ assay to assess mRNA integrity in mouse synaptosomal preparations that lack rRNAs. We concluded that the 5’:3’ assay can be used as a reliable and sensitive method to evaluate mRNA integrity in mouse and human brain tissue and subcellular preparations.

## Introduction

Good RNA quality is essential for acquiring reliable and replicable data from any gene expression analysis^1^. Purified RNA is quite sensitive to environmental factors and is prone to the degradation due to its single-stranded structure and abundant RNases in the environment. Therefore, measuring RNA integrity is established as a ubiquitous step to check RNA quality check before any further downstream analysis. First methods to test RNA integrity were based on RNA electrophoresis, where some of RNA sample was run on denaturing agarose gel, and RNA integrity was evaluated by comparing the intensity of visible small 18S and large 28S ribosomal RNA (rRNA) bands^2^. However, this method required at least a few hundred nanograms of RNA, was difficult to standardize and the conclusions heavily depended on subjective visual interpretation. Since 2006, microfluidic capillary electrophoresis became a gold standard for RNA integrity measurements assigning an RNA integrity number (RIN) to a assayed RNA sample^3^. However, both RNA gel and capillary electrophoresis predominantly evaluate rRNA integrity, assuming that it reflects the integrity of all other sample RNAs, including messenger RNA (mRNA).

Meanwhile, gene expression studies are mainly concerned with the quality of mRNA and not rRNA. It has been shown that because of structural and functional differences between them rRNA cannot represent the integrity of mRNA accurately enough^1, 4, 5^. In the cells, rRNAs are compact and contain many post-transcriptional modifications to form complex and functional ribosomes, which grants them exceptional stability^6^. In contrast, mRNA has a more linear structure and is more prone to degradation^7^. Therefore, RNases and any environmental factors such as temperature affect the stability and degradation rates of rRNA and mRNA differently^5^. Moreover, rRNA-based methods cannot be used to evaluate RNA integrity in subcellular extracts or purified organelles, where rRNA content is reduced or they are removed completely.

The integrity of mRNA in a RNA samples can be measured directly using real-time quantitative reverse transcription PCR (RT-qPCR). This approach, called 3’:5’ assay, has been first proposed in 2006 by Nolan and colleagues^8^. The 3’:5’ assay measures the integrity of the ubiquitously expressed mRNA of choice, independently of rRNAs. By now a few versions of the 3’:5’ assay have been published that used different template mRNAs for different species^8–11^. However, 3’:5’ assays accuracy has not been sufficiently explored as the few existing studies validate the assay only by comparing 3’:5’ assays integrity values to RIN values. Also, no such assay for mice or human RNA samples was developed. Furthermore, recent studies demonstrated that even minor differences of primer efficiencies result in significant under- or overestimation of mRNA levels, which should be accounted for in a qPCR based assay^12^.

Therefore, to have a convenient and reliable method for direct mRNA integrity evaluation we developed and optimized a *PGK1* transcript-based 5’:3’ assay for human and mice brain tissue. To adjust for any inevitable differences in the efficiencies of qPCR primer pairs used in the assay we refined the integrity calculation by using efficiency correction. Moreover, we aligned 5’:3’ integrity value calculation to other existing RNA integrity evaluation systems, where integrity values close to 10 indicate very good quality samples and decrease when RNA is degraded. We demonstrated that the 5’:3’ assay can be confidently used as an alternative method to measure RNA integrity in mouse and human brain tissue samples. Finally, we showed that this approach can be successfully used to evaluate RNA integrity in subcellular fractions that lack rRNA. As a model of such fraction we used synaptosomes, which are isolated synaptic terminals from the neurons. Synaptosomes are scarce of rRNA and therefore cannot be assessed by conventional rRNA-based RNA integrity methods.

## Results

### 5’:3’ assay based on *PGK1* cDNA as a template

The 5′:3′ approach is based on using oligo-dT primers for reverse transcription and two sets of primers for qPCR to measure the relative expression of two amplicons located on the 3′ and 5′ regions of a long constitutively expressed mRNA of choice and then comparing the relative amounts of 3’- and 5’-ends of the transcript (**Figure 1**). If mRNA in the sample is intact, reverse transcription from poly(A) tails goes on uninterrupted generating full-length cDNA. Therefore, following qPCR generates similar levels of 3′- and 5′-end products. In a partly degraded mRNA sample, cDNA synthesis from the poly(A) tail of fragmented mRNA leads to the truncated cDNA at the site of mRNA cleavage. This truncation depletes the binding sites for 5’-end primer pair resulting in a reduced amount of 5’-end product. The 5’:3’ integrity value for a sample is then quantified by dividing the amount of 5’-end amplicon amount by that of 3’-end amplicon and multiplying by 10 to get an integrity score from 10 (intact mRNA) to 0 (totally degraded mRNA), in line with other RNA integrity assays that use 1–10 scales^3^.

**Figure 1.**
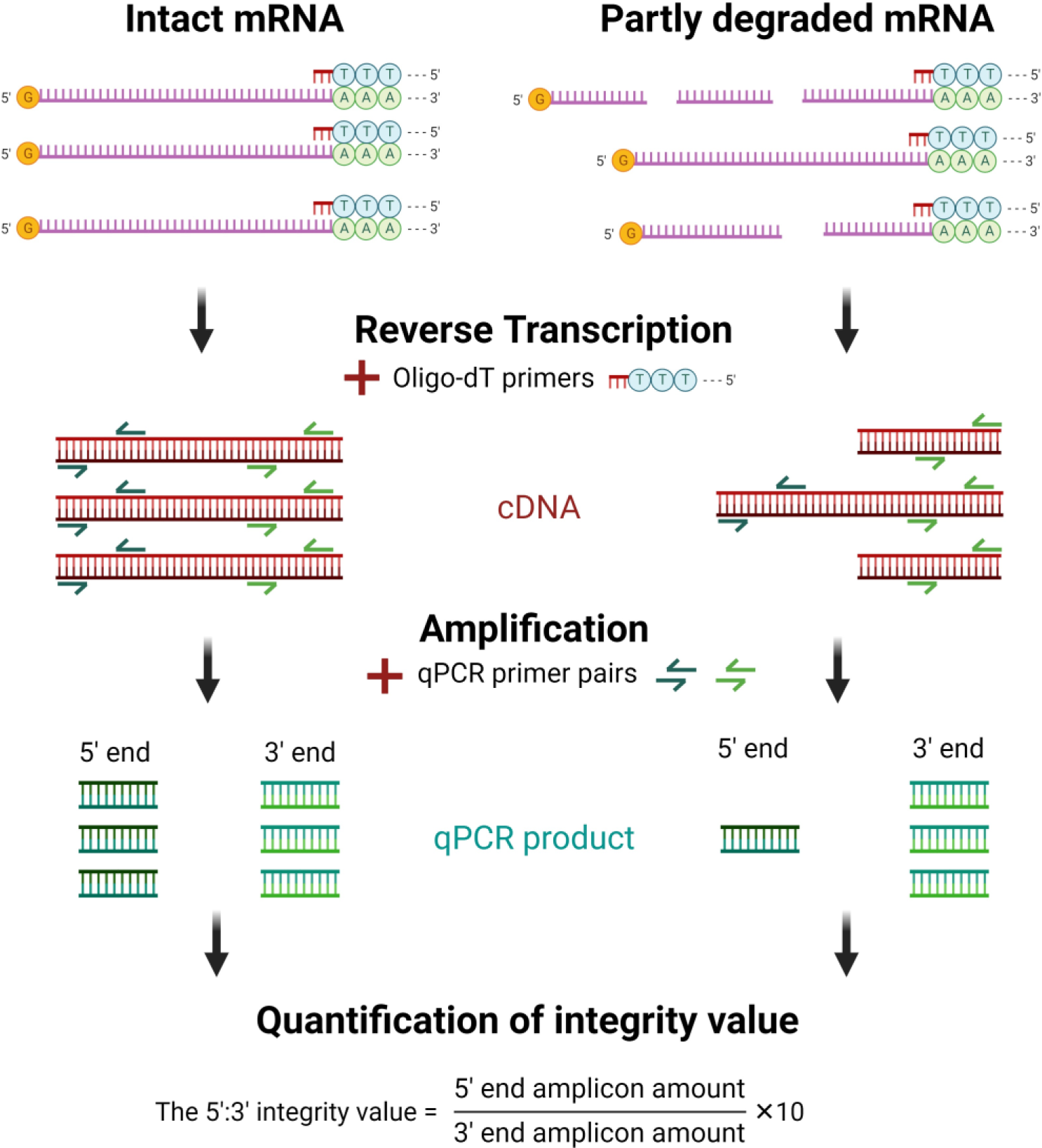
The principle of 5′:3′ assay for mRNA integrity evaluation. Reverse transcription reaction used oligo-dT primers to generate cDNA. In RNA samples of good quality, mRNA is intact and full-length cDNA is synthesized. In this case, both 3’and 5’ pairs of qPCR primers have the same number of sites to attach and produce comparable amounts of 3’- and 5’-end PCR products. In contrast, in poor quality RNA samples with partly degraded mRNA, shorter cDNA is synthesized because of the breaks in mRNA. On truncated cDNA, PCR primer pair for the 5’-end of mRNA has lower number of binding sites compared to the 3’-end, close to the poly(A) tail of mRNA. After qPCR amplification, the 5’:3’ integrity value is calculated by dividing the amount of the 5’-end amplicon by that of the 3’-end amplicon and multiplying by 10, allowing the quantitative evaluation of mRNA quality in the sample. Created with Biorender.com.

The mRNA template for 5’:3’ assay has to be carefully selected and ideally should be a long transcript with a stable and abundant expression in the tissue of interest with a few pseudogenes to ensure reliable results. Here we used mouse and human *PGK1* cDNA as a template for the 5’:3’ assay. *PGK1* is ubiquitously expressed housekeeping gene that produces a relatively long transcript with a well-characterized exon-intron structure well suited for this assay^13^. Its product is an enzyme called phosphoglycerate kinase, which is involved in glycolysis^14^. Due to its high and stable expression, *PGK1* has been recommended as a reference gene for quantitative measurements of gene expression in RNA samples isolated from human blood^15^. Moreover, *PGK1* gene is highly conserved and homologous between rats, mice and humans. In mice and humans, there are two *PGK* isozymes encoded by different genes: *PGK1* and *PGK2* but only *PGK1* is expressed ubiquitously^16^. *PGK1* transcripts have been previously used to measure mRNA integrity in rat toxicology studies^10^ and horse samples^11^. Here we designed the qPCR primers for 5’:3’ assay based on mouse and human *PGK1* cDNA to measure mRNA integrity in mouse and human RNA samples (**Table 1**).

**Table 1.**
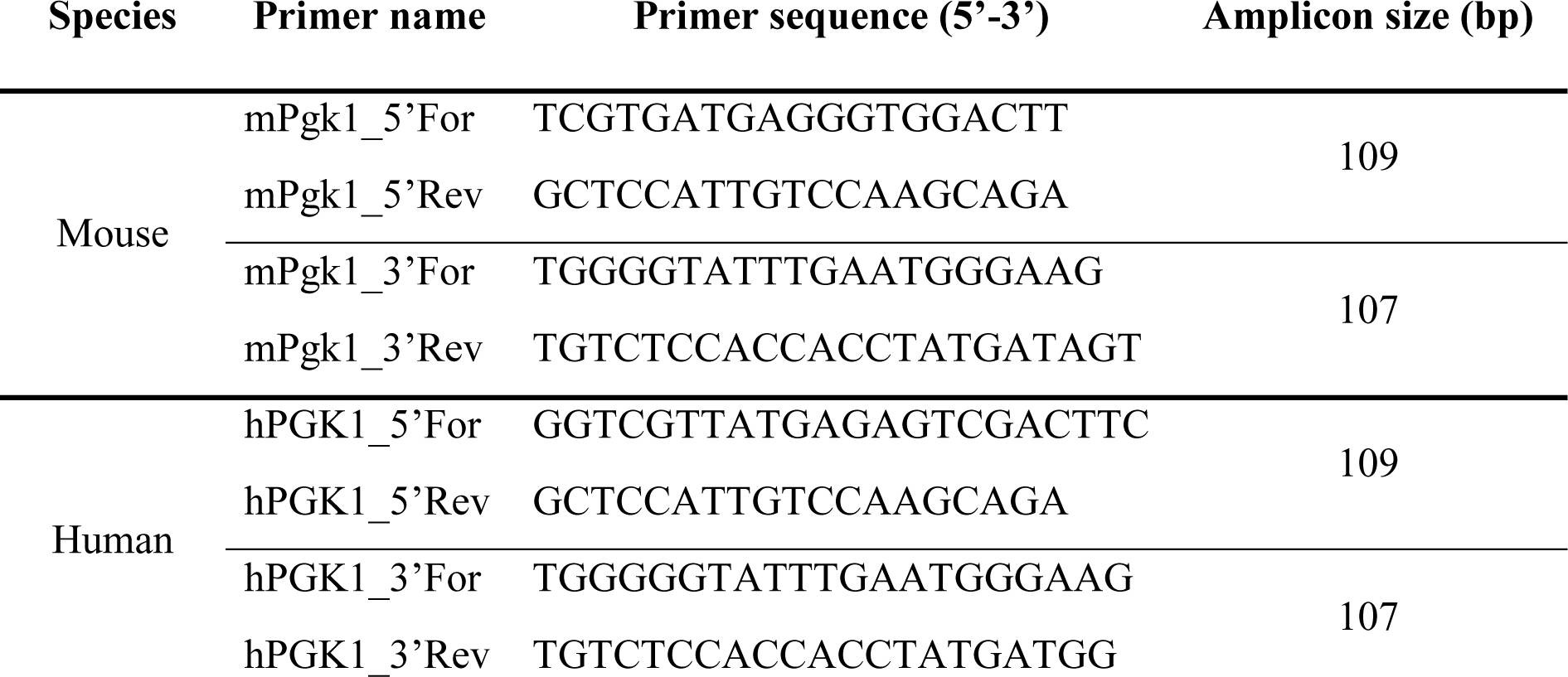
Primer pair sequences for 5’:3’ assay in mouse and human RNA samples.

### The integrity value of 5’:3’ assay is corrected to reflect primer pair amplification efficiency

The 5’:3’ integrity assay relies heavily on the prerequisite that both 5’- and 3’-end fragments are amplified with equal efficiency. Therefore, it is important to assess the amplification efficiency of used primer pairs. To calculate the amplification efficiency of designed primer pairs (**Table 1**) we used plasmids with cloned fragments of mice (55-1342 bp) and human *PGK1* (21-1270 bp) cDNA as a template for qPCR (**Supplementary Figure S1A, C**). Acquired Ct values of both 5’- and 3’-end amplicons were plotted against corresponding plasmid copy number and amplification efficiency (*E*) was calculated from the slope of linear regression equation. For both mouse and human *PGK1*, the primer pair for the 5’-end had lower amplification efficiency (**Figure 2A, B**). The same pattern was observed when amplification efficiency was assessed in total RNA samples from mouse and human brain (**Figure 2C, D**).

**Fig. 2.**
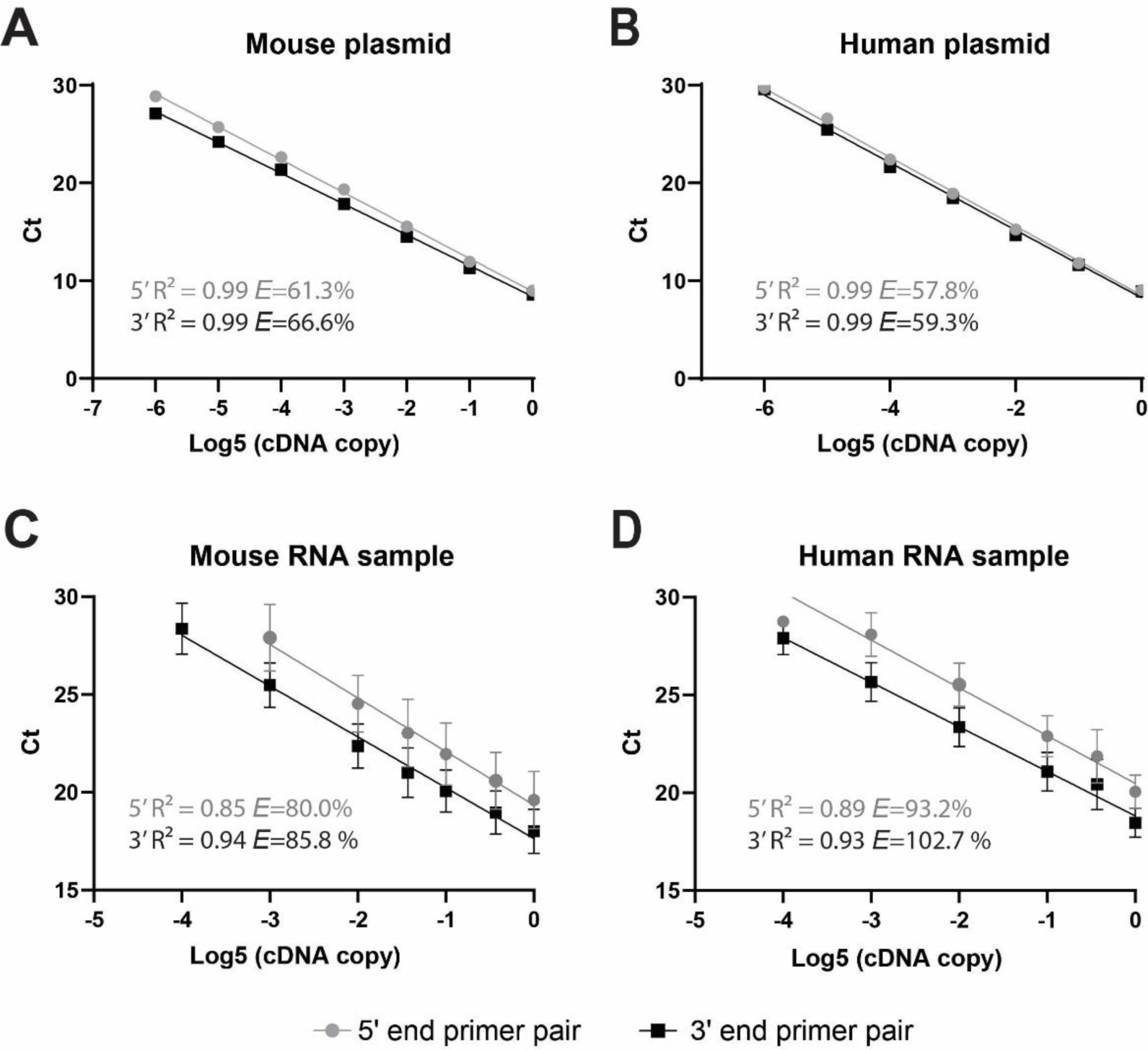
The amplification efficiency of the 5’-end and 3’-end primer pairs differs for both mouse and human 5’:3’ assay. Plasmids with cloned mouse (**A**) and human *PGK1* (**B**) cDNA or cDNA from mouse (**C,** n=2) and human (**D,** n=5) RNA samples were used for serial dilutions and following qPCR to determine primer amplification efficiency (*E*).

Even a slight difference in the primer amplification efficiency between the primer pairs results in shifted integrity values of 5’:3’ assay. Therefore, we applied a recently published STAR protocol for qPCR data analysis that corrects the relative expression of the gene(s) of interest according to primer amplification efficiency^12^. This protocol uses mathematical transformation to calculate amplification factor from the amplification efficiency (slope) for each primer pair that can be expressed as presented in the equation (*1*). Amplification factor is then used to correct the qPCR data, by defining a linear form of original gene expression (*2*). We integrated this approach to obtain adjusted 5′:3′ integrity values. To assess the outcome of this correction, we compared uncorrected and corrected 5’:3’integrity values to RIN defined in the same set of RNA samples, isolated from surgically resected human brain tissue (**Supplementary Table S1**). Paired *t*-test revealed that without the correction, the integrity values were significantly lower than RIN values, while 5’:3’ integrity values after correction were comparable to RIN values (**Supplementary Table S2**).

1. *Amplification factor = Dilution factor*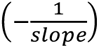
2. *Correted expression = Amplification factor^-Ct^*

### The integrity value of 5’:3’ assay accurately represents different ratios of primer binding sites

To demonstrate that the calculated integrity value translates to the actual abundance of 3’ and 5’ binding sites, we cloned plasmids with truncated mouse and human *PGK1* cDNA. These plasmids contained 3’ half of the cDNA (675-1342 bp of mouse *Pgk1* cDNA and 608-1270 bp of human *PGK1* cDNA) (**Supplementary Figure S1B, D**). By mixing different quantities of plasmids with cloned full-length *PGK1* and truncated 3’-end *PGK1* fragment, we prepared samples with known 3’- and 5’-end ratios to model variable cDNA sets resulting from RNA samples degraded to different extent. The 5’:3’ assay of these samples demonstrated a linear relationship between 3’- and 5’-end ratios and obtained integrity value (**Figure 3**). Linear relationship was also observed in the samples representing low degradation spectrum: from non-degraded sample having equal amount of 5’-end and 3’-end sites to the samples, in which the amount of 5’-end sites was reduced by half (**Figure 3, *zoomed in plots***). The linear regression lines did not differ between the wide (3’- to 5’-ends ratio from 1 to 2^6) and narrow (3’- to 5’- ends ratio from 1 to 2) intervals (*p* = 0.46 and *p* = 0.44 for mouse samples; *p* = 0.45 and *p* = 0.98 for human samples; slopes and intercepts, respectively), indicating a stable linear relationship through the whole tested range of 5’- and 3’-ends ratios and integrity values. These results showed that 5’:3’ assays integrity value can accurately represent the difference in the abundance of existing binding sites for created primer pairs.

**Figure 3.**
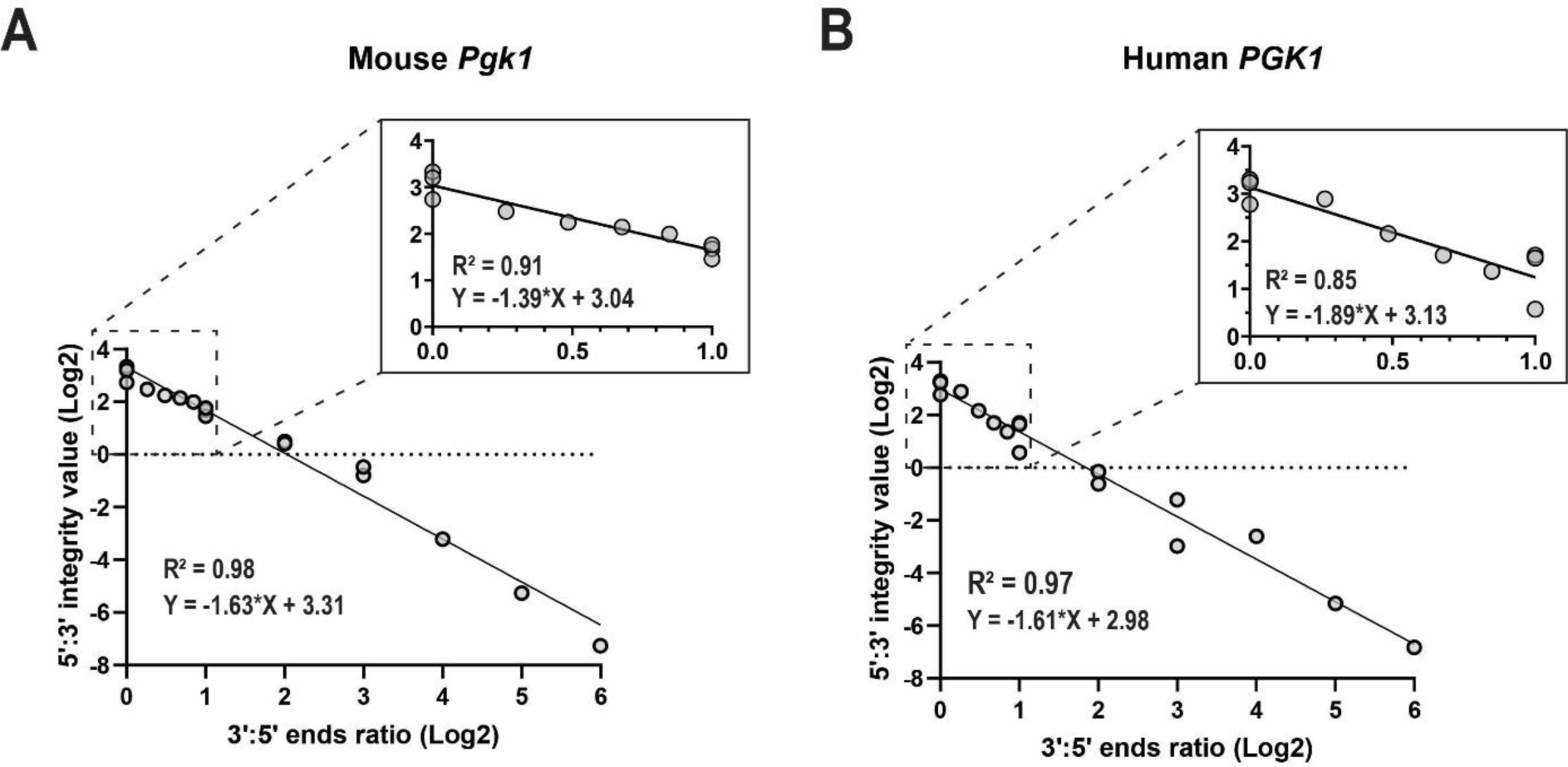
The calculated 5‘:3‘ integrity value well represents the ratios of the binding sites for the used primer pairs. The different 5’:3’ ends ratios were acquired by mixing plasmids with cloned full length and truncated mouse and human *PGK1* cDNA to mimic a wide range of mRNA degradation. The 5‘:3‘ integrity value was calculated and plotted against the ends ratio for (**A)** mouse and (**B)** human assays. The smaller graphs in (**A**) and (**B**) are zoomed in plots representing PGK1 gene 5’- and 3’-ends ratios from 1 to 2 and simulating low mRNA degradation. A stable linear relationship was observed between 5’:3’ends ratios and integrity values for both mouse and human assays.

### 5’:3’ integrity value and RIN value correlate well in heat treated RNA samples

Traditionally, RNA integrity is evaluated qualitatively by inspecting the intensities of the 28S and 18S ribosomal RNA (rRNA). This principle is used by currently most popular approach to assess RNA integrity – The Agilent Bioanalyzer system, which assigns RNA Integrity Number (RIN) to RNA sample. To assess the capacity of the 5’:3’assay to detect the degradation in general RNA samples compared to RIN value, we used heat-degraded total RNA samples from mouse cortex. The samples were incubated for 1–15 min at 90 °C and then analysed using mouse 5′:3′ RNA integrity assay and Agilent Bioanalyzer. Both the 5′:3′ integrity values and RIN values gradually decreased with prolonged heat treatment (**Figure 4A, C, D**). RIN values and 5′:3′ integrity values correlated well (R^2^ = 0.67, *p* < 0.0001) and samples that had lower RIN value had lower 5′:3′ integrity value and *vice versa* (**Figure 4B**). In primary samples, the values calculated with both methods were close (8.3 and 8.56 for RIN and 5′:3′ integrity value, respectively), but with prolonged heating 5′:3′ assays integrity values dropped faster than RIN values (**Figure 4A, C, D**). Therefore, RIN cut-of value of 7.0, which is commonly used as a cut-off for good quality RNA samples suitable for further analysis, is equivalent to 5′:3′ assays integrity value of approximately 6.

**Fig. 4.**
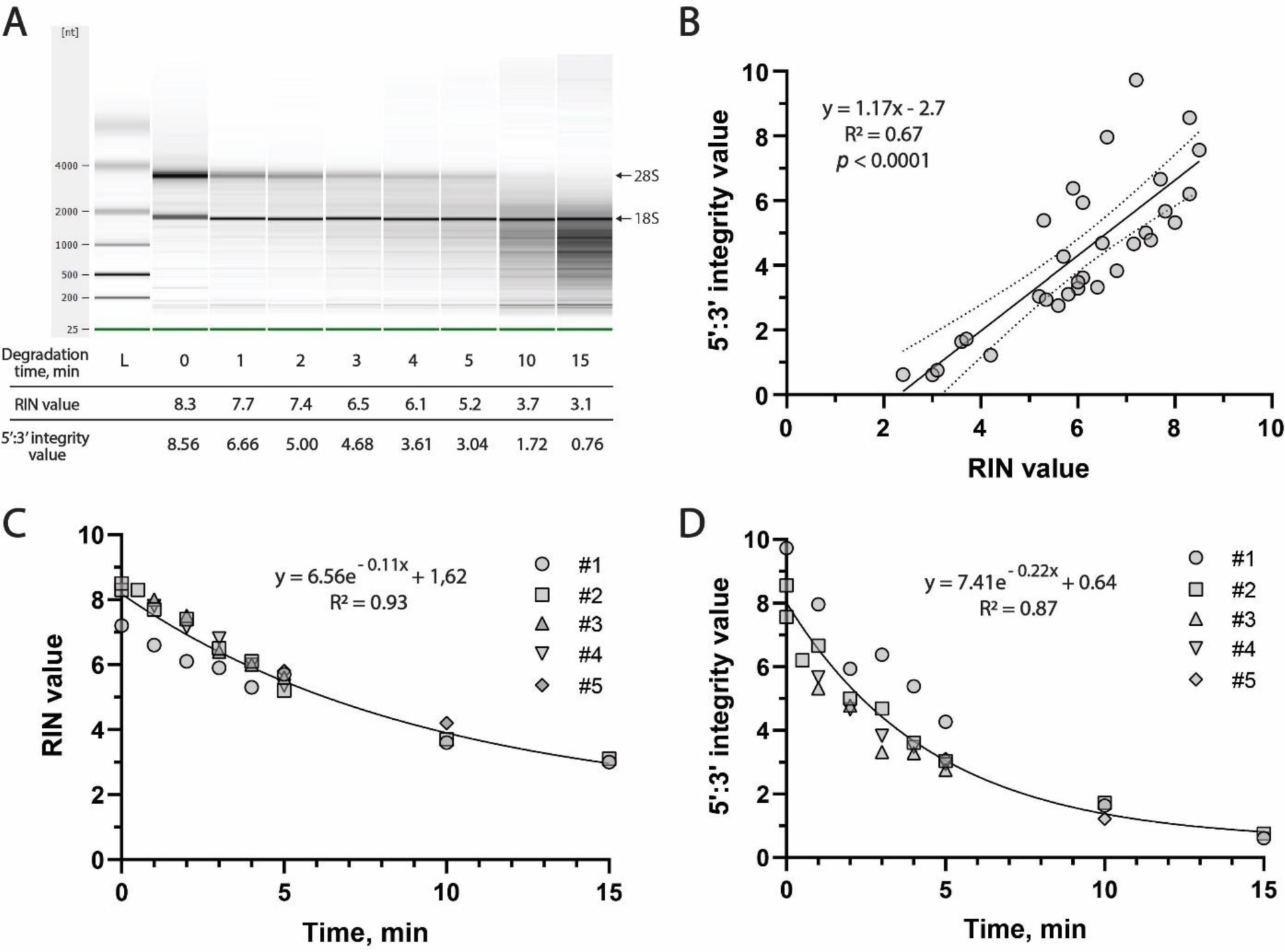
Evaluating RNA integrity with 5‘:3‘ assay *versus* RIN values using heat-degraded mouse RNA samples. **A** Representative image of Agilent 2100 Bioanalyzer RNA Pico Chip gel of mouse brain total RNA samples heat-degraded at 90 °C for 1, 2, 3, 4, 5, 10 and 15 min. At the bottom – RIN values and obtained 5’:3’ values. **B** There was a strong linear correlation between RIN value and 5’:3’ integrity value (n=5). Linear regression line is shown with 95% confidence interval of the best-fit line. **C, D** Both RIN values (**C**) and 5’:3’ integrity values (**D**) decreased with longer heating duration. The lines in **C** and **D** represents phase exponential decay of the data.

Next, we applied the same principle to assess 5’:3′ RNA integrity assay with human brain RNA samples. Three total RNA samples from surgically resected human brain tissue were heat-degraded for 1–10 min at 90 °C and then analyzed using the Agilent Bioanalyzer and human 5′:3′ RNA integrity assay. As seen with mouse RNA samples, both 5’:3′ integrity values and RIN values decreased with longer duration of sample heating (**Figure 5A, C, D**). There was a very strong correlation between 5′:3′ RNA integrity values and RIN values (R^2^ = 0.86, *p* < 0.0001) showing that both values reflect RNA degradation in samples comparatively (**Figure 5B**). As with mouse RNA samples, heat degradation decreased human 5′:3′ integrity values faster than RIN values (**Figure 5A, C, D**).

**Figure 5.**
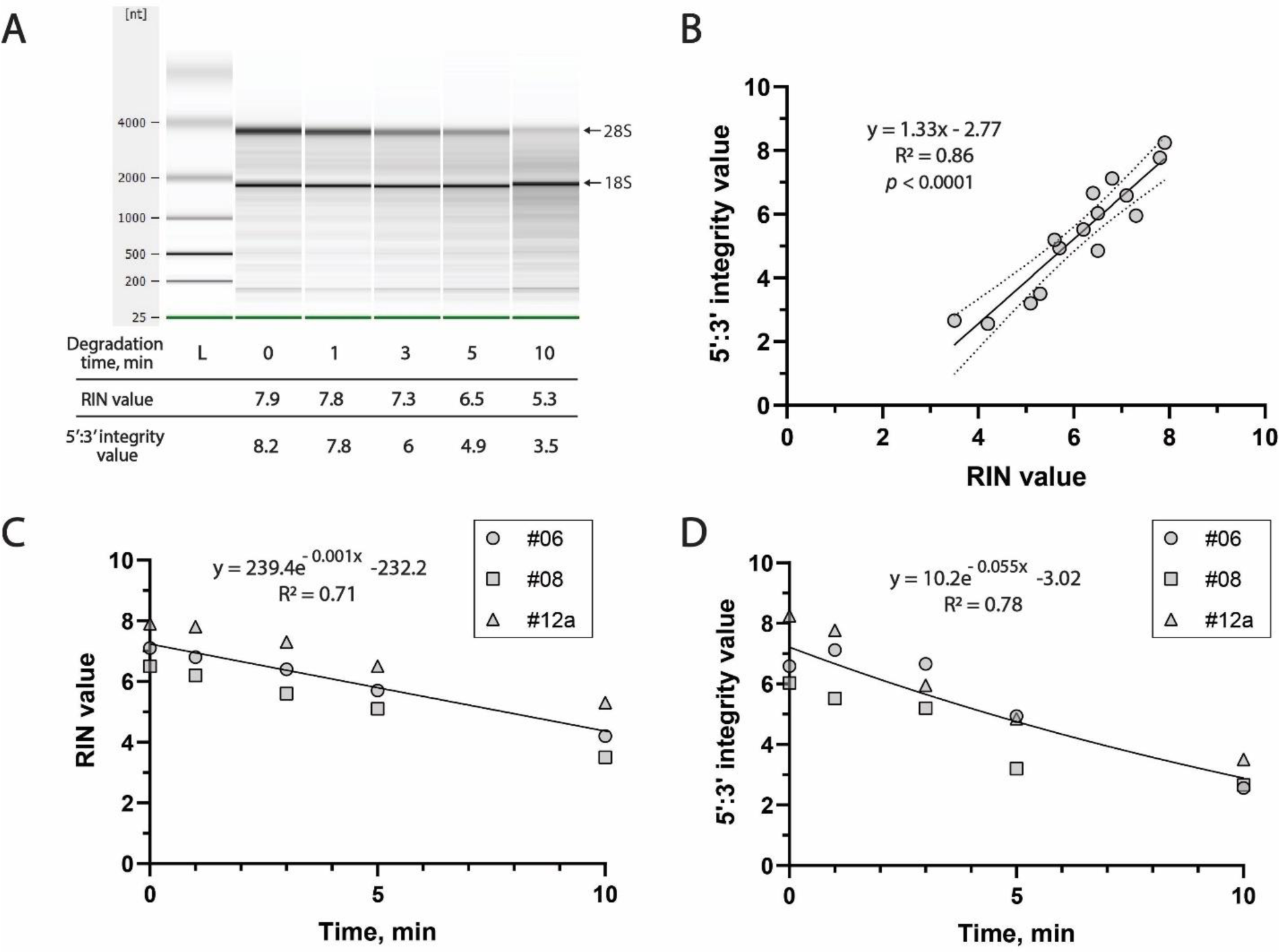
Evaluating RNA integrity with human 5‘:3‘ assay *versus* RIN values using heat-degraded human brain RNA samples. **A** Representative image of Agilent 2100 Bioanalyzer RNA Pico Chip gel of human brain total RNA samples heat-degraded at 90 °C for 1, 3, 5, and 10 min. At the bottom – RIN values and obtained 5’:3’ values. **B** Three total RNA samples from surgically resected human brain tissue were used to define 5’:3’ integrity values and RIN values (#06, #08, #12a, **Supplementary Table S1**). There was a strong linear correlation between RIN values and 5’:3’ integrity values in these samples. Linear regression line is shown with 95% confidence bands of the best-fit line. **C, D** Both RIN values (C) and 5’:3’ integrity values (D) decreased with longer heating duration. The lines in **C** and **D** represent one phase exponential decay of the data.

### In human post-surgical brain tissue samples, 5‘:3‘ integrity value represents the RNA integrity better than RIN

RT-qPCR based RNA integrity evaluation method can be applied as an RNA quality control method for most medical laboratories working with human tissues. The increasing use of human tissue in research emphasize the importance of developing appropriate and easy to use methods for tissue quality assessment. For that reason, we assessed human 5’:3’ assay to evaluate RNA integrity in human brain samples, obtained from the glioma or epilepsy surgery. RIN values were measured in the same samples for comparison. The surgically resected brain tissue was kept viable during transportation in cold artificial Cerebrospinal Fluid (aCSF) and then dissected immediately for homogenization, RNA extraction and both RNA integrity assays (**Figure 6A**). Obtained 5’:3’ integrity values varied from 3.6 to 9.7 while RIN values varied from 4.6 to 7.8 in these samples. There was a significant moderate correlation between 5’:3’ integrity assay and RIN values (R^2^ = 0.59, *p* = 0.0005) (**Figure 6B**). Therefore, 5’:3’ integrity value can be used to reliably evaluate RNA quality in clinical samples as an alternative to RIN.

**Fig. 6.**
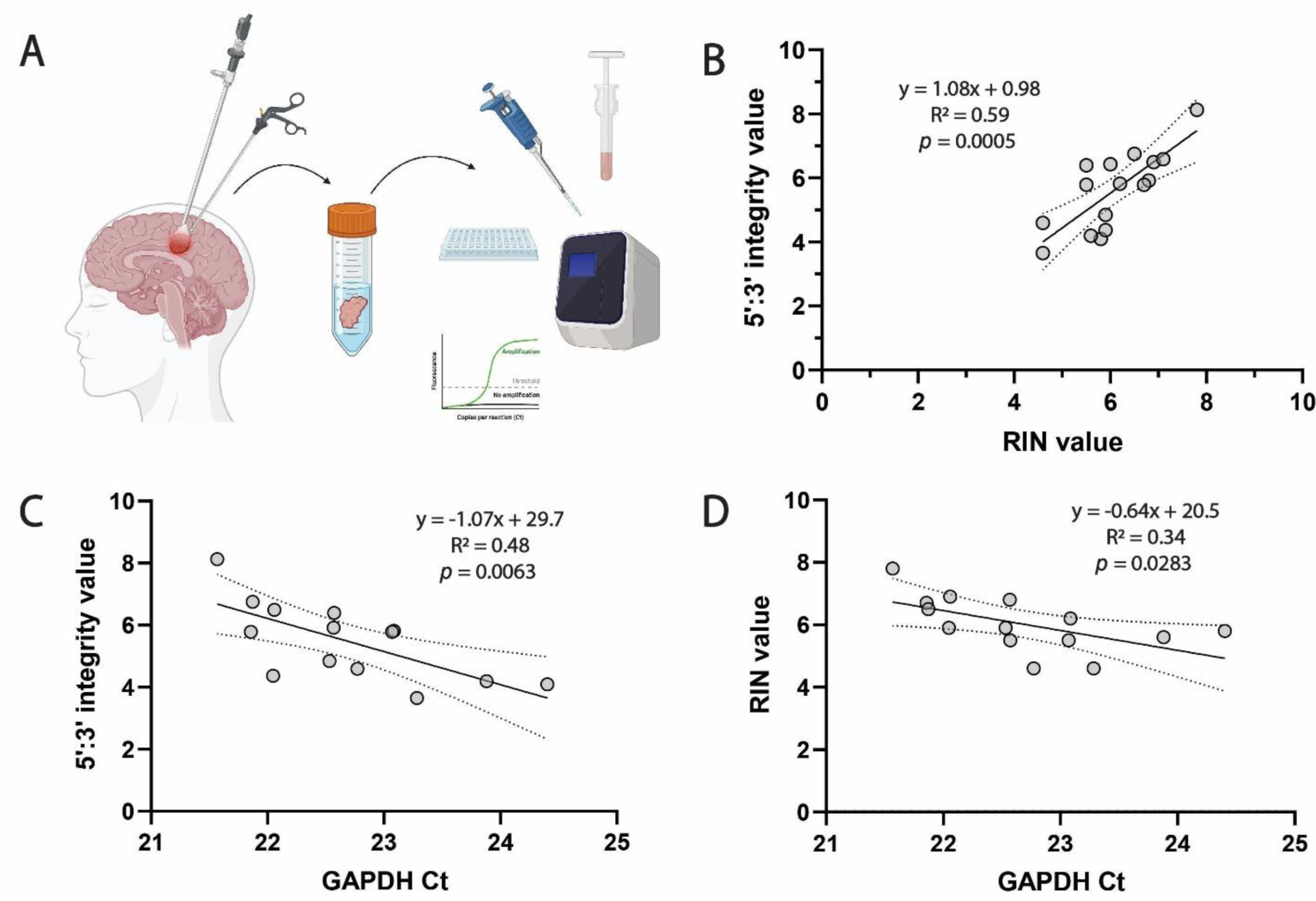
RNA integrity in total RNA samples from surgically resected human brain tissue. **A** Human post-surgical brain tissue from the glioma or epilepsy surgery was kept viable during transportation in artificial cerebrospinal fluid (aCSF). The sample was then immediately dissected and homogenized for RNA extraction and RNA integrity assays. Created with Biorender.com. **B** Relationship between 5’:3’ integrity value and RIN in RNA samples from human brain tissue (n=16). Linear regression line in **B** is shown with 95% confidence intervals of the best fit line. **C** There was a moderate linear correlation between the human brain tissue RNA sample 5’:3’ integrity values and Ct values from qPCR for *GAPDH* expression, while the correlation with RIN values was weak (**D**) (n=14).

For gene expression studies, it is important to define the integrity of mRNA in particular rather than total RNA or rRNA, which is defined by RIN. It is known that RNA integrity determines acquired Ct values in qPCR reactions for gene expression studies^1^. RNA degradation lowers the abundance of the binding sites for qPCR primers and leads to higher Ct values compared to intact RNA samples. Therefore, we compared 5’:3’ integrity values and RIN values to the expression of a housekeeping gene *GAPDH* in the same clinical samples as defined by real-time qPCR using TaqMan system. The 5’:3’ integrity value correlated better with *GAPDH* Ct values then RIN values, which further supported our assertion that 5’:3’assay reflected transcript quality in RNA samples better than RIN value (R^2^ = 0.48, *p* = 0.0063 for 5’:3’ integrity value and R^2^ = 0.34, *p* = 0.0283 for RIN value) (**Figure 6C, D**). Importantly, when another highly expressed *CMAS* gene was used for comparison, only 5’:3’ integrity value correlated significantly with the results of defined expression of *CMAS* transcripts (**Supplementary Figure S2**), confirming that direct evaluation of mRNA integrity is more reliable than other methods for RNA quality assessment.

### 3′:5′ assay can be used to measure RNA integrity in synaptosomal preparations

Measuring RNA integrity in subcellular fraction samples presents further challenges, as such samples usually lack sufficient quantities of ribosomal RNA to determine RIN values. The examples of such fractions are various extracellular vesicles or synaptosomes (isolated neuronal synaptic terminals). Pure synaptosome sample can be obtained by using genetically engineered mouse where synapses are fluorescently labelled together with fluorescence activated synaptosome sorting (FASS) to sort and collect the labelled synapses. It has been shown that because of its ubiquitous expression *PGK1* is abundant in synaptosomes^17^. Therefore, to test whether 5’:3’ integrity assay can be used to evaluate RNA integrity in subcellular fractions, as a model, we used RNA from eight matched synaptosomal preparations – four crude synaptosome samples and four FASS-enriched synaptosome samples. Crude synaptosome samples were prepared by gently homogenizing vGLUT1^mVenus^ mouse^18^ cortical tissue and fractionating synaptosomes in sucrose gradient. To enrich the samples for excitatory synaptosomes, we used FASS technique to sort out mVenus^+^ synaptosomes from crude synaptosomal sample^19^ (**Figure 7A**). After the sorting, 60-90 % of all the particles were mVenus+ compared to ∼20% in crude preparations, indicating significant excitatory synapse enrichment (**Supplementary Figure S3**). Crude synaptosome samples were a mixture of similar size and density particles still containing some rRNA. Therefore, RNA integrity analysis in crude synaptosome samples demonstrated comparable values for 5’:3’ integrity assay and RIN. However, in the enriched synaptosomal samples there was almost no ribosomal RNA, thus there were no detected rRNA bands in electrophoresis lanes and RIN values could not be determined. In contrast, 3′:5′ assay, which is independent of rRNA, was suitable to evaluate mRNA integrity in RNA samples from enriched synaptosomes (**Figure 7B, C**), demonstrating its applicability for the samples containing subcellular fractions.

**Figure 7.**
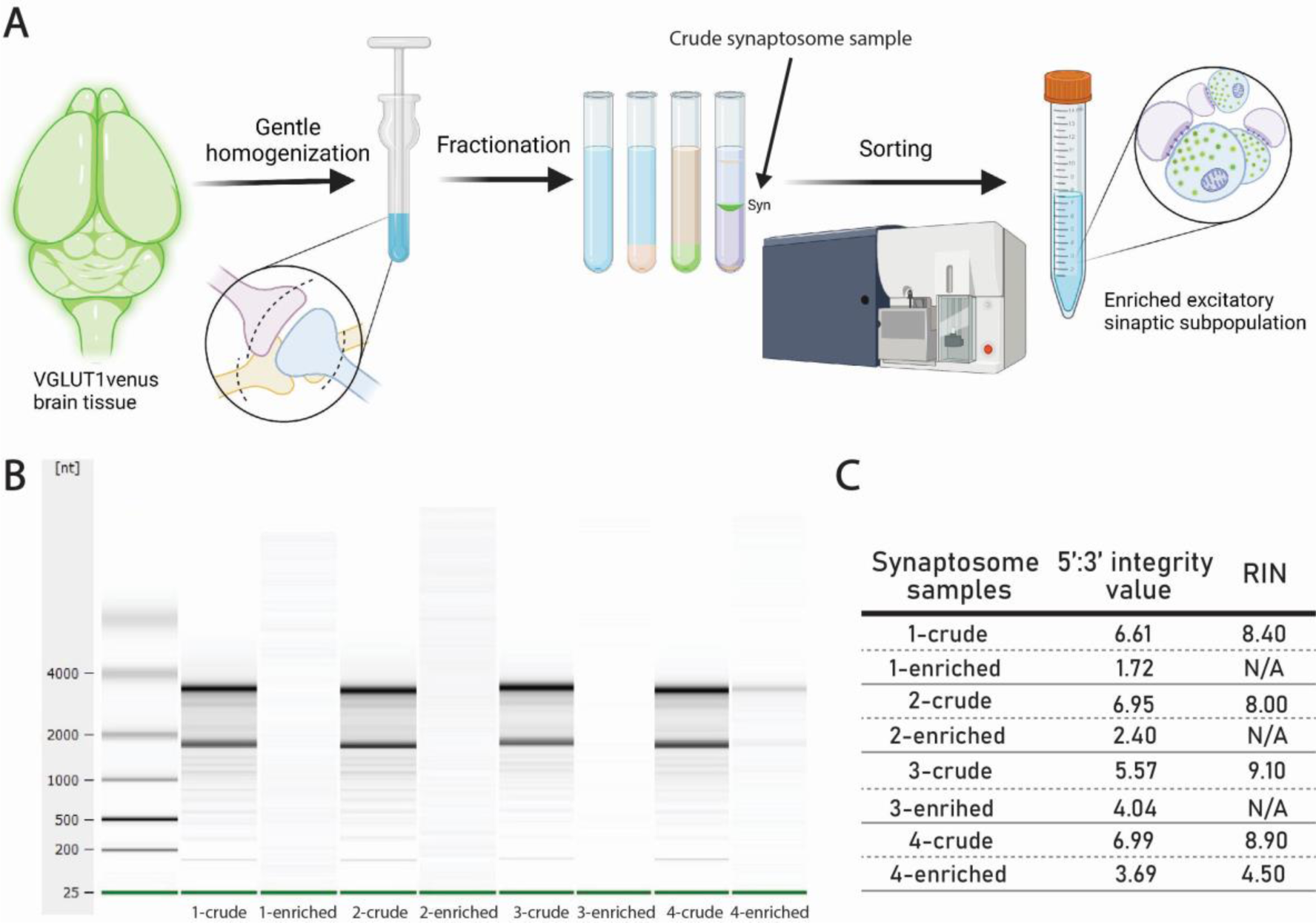
Evaluating RNA integrity in synaptosomes. **A** VGLUT1^mVenus^ mouse brain tissue was homogenized and the homogenate was serially fractionated to obtain the particles that have the size and the density corresponding to the synaptosomes. This crude synaptosome sample was then sorted according to mVenus fluorescence to get enriched excitatory synaptosome subpopulation. Created with Biorender.com. **B** Representative image of Agilent 2100 Bioanalyzer RNA Pico Chip gel of total RNA samples from crude and enriched mouse synaptosomal samples. **C** 5’:3’ integrity value and RIN value of crude and enriched mouse synaptosomal samples.

## Discussion

In healthy cells, RNA degradation is strictly regulated^7^. However, during sample collection this homeostatic regulation is disrupted and RNA degradation is enhanced. The degradation of RNA samples undermines the accuracy of gene expression analysis by qPCR or RNA sequencing^1^. RNA sequencing is particularly sensitive to mRNA degradation, as mRNAs are usually captured by poly-dT probes in order to deplete overabundant rRNA. When an mRNA sample is degraded, only a few full length transcripts are captured, which leads to the 3’ bias of RNA sequencing results^20, 21^. Due to this 3’ bias, the reads are predominantly limited to the last exon of mRNA, which distorts gene expression profiles compared to not degraded RNA samples^20^. Therefore, to obtain reliable and reproducible gene expression data, it is crucial to take into account the integrity of sample mRNA. Conventionally, RNA integrity is measured by RNA electrophoresis-based methods. However, these methods rely on the assumption that rRNA integrity represents the mRNA integrity well enough, even though it has been shown not to be the case^4, 5^. Therefore, we developed a direct mRNA integrity evaluation assay that does not require any specialized equipment and is based on the comparison of the abundance of 5’:3’ transcripts of housekeeping genes in mouse and human RNA samples.

This mouse and human 5’:3’ assay is based on *PGK1* mRNA as it is a long constitutively expressed transcript of a highly conserved gene^16^. The *PGK1* RNA was previously successfully used for designing 3’:5’ assay primers for rat and horse RNA samples^10, 11^. Several other genes have also been tested in the past. The 3’:5’ assay was first published based on human *GAPDH* gene as it is a popular choice for endogenous control^8^. There was also a variant of this method based on *β-actin* mRNA optimized for sea bass larvae^9^. However, there have been reports demonstrating high expression variability of *GAPDH*^22^. It is also well known that *GAPDH* and *β-actin* have many pseudogenes^23–25^. This means that the results obtained using these genes as constitutive controls might not translate very well across different tissues or conditions and it will be more susceptible to errors caused by genomic DNA contamination.

Having an easily understandable scale for 5’:3’ assay was one of our goals to make it convenient for users. Commercial systems designed for the assessment of RNA integrity, usually give values that span from 1 (totally degraded RNA) to 10 (intact RNA). All of the previously published 3’:5’ assay protocols used the opposite scale, in which high quality RNA samples are represented by integrity values close to 1. This was unnecessarily confusing for the researchers and provided challenges when comparing different RNA integrity assays. Therefore, we established a reversed 5’:3’ integrity value to have the same scale as RIN.

Another important improvement of 5’:3’assay was the correction of the integrity score according to the amplification efficiency of used primer pairs. Primer amplification efficiency is known to fluctuate between different primers. This cannot be completely avoided during primer design process and highly depends on the qPCR machine, the choice of reagents for PCR, the presence of various inhibitors in the sample and even the sample volume used for the dilutions^26^. Therefore, it is important to correct the real-time PCR data using primer amplification efficiency to enhance the analytical accuracy^12^. We added this correction to our integrity value calculations ensuring a robust method for RNA integrity assessment. This will help to successfully establish this method in any other laboratory as the correction eliminates the bias introduced by using different primers, equipment or reagents.

To validate 5’:3’assays ability to reflect the RNA integrity status we chose to compare our results with widely used RIN value. 3‘:5‘ assay integrity value and RIN values were very comparable in heat-degraded mouse and human RNA samples (**Fig. 4b and Fig. 5b**). However, in clinical samples this correlation was lower **(Fig. 6b)** indicating that enzymatic RNA degradation, which occurs during sample collection and preparation, might affect messenger and ribosomal RNA differently. Heat-induced RNA degradation does not depend on RNA structure and thus affects both rRNA and mRNA similarly. However, mRNA is significantly more sensitive to enzymatic degradation by nuclease activity compared to rRNA^4^. In clinical samples or any other RNA samples, where RNA degradation is caused by endogenous nucleases, the results from rRNA-based methods can overestimate the RNA integrity. As sample collection in clinical settings and field studies are prone to rapid tissue necrosis, which is associated with enzymatic degradation of nucleic acids, 5’:3’ assay would be a better indicator of RNA quality in such samples.

Ribosomal RNA-based methods for RNA integrity evaluation cannot be applied for various subcellular samples, such as synaptosomes or extracellular vesicles. Such samples share similar features that affect the assessment of the expression of their local transcripts: prolonged processing of the sample, low yields of RNA and most importantly – lack of rRNA^27^. For example, we showed that extracted excitatory synaptosomes have very little rRNA (**Fig. 7b**). Therefore, RIN could not be determined in such samples, but mRNA based 5’:3’ assay was suitable to measure the RNA integrity (**Fig. 7c**). This shows a huge advantage and potential of this method to be used to check RNA integrity on any sample that lacks high quantities of rRNA, making impossible the use of common RNA integrity assays.

5’:3’ assay holds a potential as an easily applicable method with improved accuracy for estimating RNA degradation for gene expression studies. However, there are a few limitations worth mentioning. All RNA integrity evaluation strategies are based on generalized assumptions about a very diverse nucleic acid pool. For example, it’s been shown that the degree of degradation varies among different transcripts, and a substantial fraction of the variation can be explained by functional and structural features of different transcripts^20^. Also, 5’:3’ assay based on *PGK1* transcript might slightly overestimate the integrity of short transcripts as it is known that shorter transcripts degrade faster than longer ones^28^. Therefore, more research is needed to understand natural mRNA degradation process and how it affects different transcripts. Nevertheless, one of the advantages of 5’:3’ assay is that it can be easily tailored by designing new qPCR primers for a specific transcript which is then taken as a representative of mRNA pool. Thus such transcript can be chosen to represent not all but a fraction of mRNAs with certain length, structure or function of interest. Such adaptation of 5’:3’ assay to reflect a certain group of similar transcripts could improve the accuracy of mRNA integrity evaluation even further.

In conclusion, the 5’:3’ assay is a promising method for assessing mRNA integrity for gene expression studies, offering advantages over conventional techniques. The assay, based on *PGK1* mRNA, provides a user-friendly assessment system, aligning the integrity score with the widely used RIN values. Its correction for primer amplification efficiency enhances analytical accuracy, addressing the variations in qPCR conditions. While validation against RIN values showed strong correlation in heat-degraded samples, a slight disparity of correlation in clinical samples emphasizes the likely impact of the differences in enzymatic degradation of mRNA and rRNA. Despite acknowledged limitations, the adaptability of the 5’:3’ assay by designing primers for specific transcripts holds high potential for personalized assessments, offering a step forward in refining the accuracy of mRNA integrity evaluations.

## Methods

### Mouse brain tissue samples

C57BL/6J mice were obtained from local Life Sciences Center colonies. VGLUT1^mVenus^ mice^18^ were used in the homozygous state. Animal studies were conducted in accordance with the requirements of the Directive 2010/63/EU and were approved by the Lithuanian State Food and Veterinary Service (permit No. G2-92). All mice were bred and kept at the animal facility of the Life Sciences Center of Vilnius University.

For total brain RNA extraction, mouse brains were harvested from adult C57BL/6 mouse after cervical dislocation. The cortex was immediately dissected, homogenised in Trizol and stored at −80 °C until RNA isolation. VGLUT1^mVenus^ mice aged P21, P28 or P35 were used for synaptosomal preparations.

### Human brain tissue samples

Permission to use human brain tissue was obtained from Vilnius Regional Biomedical Research Ethics Committee (Approval Nr. 2020/2-1202-687). Human brain tissue specimens were obtained with informed consent as requested by the regional ethics committee (Nr.2/2020 02 18). Human neocortical access tissues (n = 16) were sampled from either glioma tumour resection surgery (n = 6), using distant cortex without tumour infiltration, or access cortex tissue from epilepsy surgery (n = 5) obtained during the resection of epileptic foci. Human hippocampus tissue (n = 5) were samples resected during epilepsy surgery. Detailed information regarding the donors is provided in **Supplementary table S1**.

### Human brain tissue preparation

Tissue was immersed in 4 °C Artificial Cerebrospinal Fluid (aCSF) immediately post-resection and transferred for homogenization in Trizol (#15596026, ThermoFisher Scientific). Homogenized samples were stored at −80 °C untill further procedures. ACSF used for tissue transport was prepared as previously reported^29^. Shortly, the solution contained 0.5 mM calcium chloride (dehydrate), 25 mM D-glucose, 20 mM HEPES, 10 mM magnesium sulfate, 1.2 mM sodium phosphate monobasic monohydrate, 92 mM N-methyl-d-glucamine chloride (NMDG-Cl), 2.5 mM potassium chloride, 30 mM sodium bicarbonate, 5 mM sodium L-ascorbate, 3 mM sodium pyruvate, and 2 mM thiourea. Prior to use, the solution was equilibrated with 95% O_2_, 5% CO_2_. Osmolality was verified to be between 295-305 mOsm/kg. pH was adjusted to 7.3 using HCl.

### Plasmids

Mouse and human *PGK1* cDNA fragments were copied and amplified by PCR using Thermo Scientific Phusion High-Fidelity DNA polymerase from mouse or human brain total RNA sample. The following primers were used: *Pgk1_For_MMPCR*: 5’-GGAGGCCCGGCATTCTGCAC-3’ and *Pgk1_Rev_MMPCR*: 5’- ACCGCCCCCAGTGCTCACATG-3’ for mouse gene and *PGK1_For_HSPCR*: 5’- GGCAGTCGGCTCCCTCGTTG-3’, *PGK1_Rev_HSPCR*: 5’- CCACCCCCAGTGCTCACATG-3’ for human gene. The PCR fragments were then ligated into pJET1.2/blunt plasmid using CloneJET PCR cloning Kit (Thermo Scientific). To generate plasmids containing 3’-end of *PGK1* cDNA 5’ part of *PGK1* was excised by NcoI from previously generated full-length *PGK1* cDNA plasmids. All pasmids were linearized by XhoI for experiments.

### Crude synaptosome sample preparation

Fresh mouse brain was cleaned and cooled in ice cold PBS. All following steps were carried on ice. Mouse visual cortex was dissected and placed into a 2 cm clean ice-cold glass-Teflon potter. The tissue was then homogenized in 1 ml of ice cold 0.32 M sucrose buffer (0.32 M sucrose, 4 mM HEPES, 4 U/μL Ribolock, pH 7.4) at 900 rpm for 15s moving the pestle up and down. The homogenate was centrifuged at 1000×g for 5 min at 4 °C. The supernatant was then centrifuged at 12,500×g for 8 min at 4 °C. The second supernatant was discarded and the pellet was carefully resuspended in ice-cold 0.3 ml 0.32 M sucrose buffer. Discontinuous sucrose gradient was prepared in 5 ml ultracentrifuge tubes (Beckman Coulter, C14279) with 2 ml of 1.2 M sucrose solution (1.2 M sucrose, 4 mM HEPES, 0.2 U/μL Ribolock, pH 7.4) at the bottom and 2ml 0.8 M sucrose solution (0.8 M sucrose, 4 mM HEPES, 0.2 U/μL Ribolock, pH 7.4) at the top. The resuspended pellet was then carefully layed over the prepared gradient. The tubes were ultracentrifuged at 24,000 rpm for 45 min at 4 °C. After ultracentrifugation two layers of particles and a pellet appeared. The middle layer containing synaptic particles was collected with a syringe by piercing through the tube wall. Collected crude synaptosome sample was then diluted in 8 ml of ice-cold 0.22 μm filtered PBS. This sample was then split and part of it was collected onto a 0.1 μm polycarbonate filter and washed with Trizol reagent for RNA extraction. Second part was used for Fluorescence Activated Synaptosome Sorting (FASS) to prepare matched excitatory synaptosome-enriched samples.

### Fluorescence Activated Synaptosome Sorting (FASS)

The prepared crude synaptosome sample was diluted in PBS and stained with SynaptoRed C2 (1μg/ml, Tocris Bioscience) to label all plasma membrane-containing particles to increase synaptosome detection sensitivity. The dilution was optimized every time to reach around 15 000 events per second at the flow rate between 3 and 4. New sorting sample was prepared every 45 min and the collection tube was changed at the same time. Synaptosome sorting was performed as described previously^19^. For sorting we used BD FACSAria III Cell Sorter set as follows: 70 μm Nozzle, sample shaking at 300 rpm, sample temperature at 4 °C, FSC neutral density filter 1.0, 488 nm Laser on, Area Scaling 1.18, Window Extension 0.0, FSC Area Scaling 0.96, Sort Precision: 0-16-0. Thresholding on SynaptoRed C2 with a threshold value of 800. Synaptosomes were sorted into a 5 ml polystyrene round-bottom tubes (BD Falcon) and kept on ice. The synaptosomes were sorted and collected for at least 6 hours. Then synaptosomes were collected onto a 0.1 μm polycarbonate filter and washed with Trizol reagent for RNA extraction.

### RNA extraction

Total RNA from mouse cortex, crude and enriched synaptosomal preparations and surgically resected human brain samples was extracted using TRIzol Plus RNA Purification Kit and Phasemaker Tubes Complete System (Invitrogen, cat. no. A33254). The remaining gDNA was removed using TURBO DNA-free Kit (Invitrogen, cat. no. AM1907) following the manufacturer’s protocols.

### Preparation of heat-degraded RNA samples

Mouse cortex RNA samples were diluted to 100 ng/μl. An aliquot (12 μl) of each diluted RNA sample was taken into separate tubes. While one tube served as the control (heated at 70 °C for 2 min to denature RNA), the other tubes were exposed to 90 °C heat for 1, 2, 3, 4, 5, 10, or 15 min in a thermocycler. Same procedure was applied to human brain total RNA samples which were kept at 90 °C for 1, 3, 5, or 10 min.

### Evaluation of RNA concentration and rRNA integrity

A NanoDrop 2000 Spectrophotometer (ThermoFisher Scientific, Waltham, MA USA) was used to measure the absorbance at 260 nm to evaluate RNA concentration. rRNA integrity was assessed by an Agilent 2100 Bioanalyzer system, using the Agilent RNA 6000 Pico Kit (Part Number: 5067-1513) (Agilent Technologies, Mississauga, ON, Canada) following the manufacturer’s protocols.

### cDNA synthesis and RT-qPCR

The reverse transcription reactions were performed in a total volume of 20 μl using 1 μg of total RNA (heated at 70 ᵒC for 2 min to denature RNA), 2.5 μg Anchored Oligo-dT primers (Invitrogen), and 200 U Thermo Scientific Maxima Reverse Transcriptase (Thermo Scientific) following manufacturer’s first-strand cDNA synthesis protocol. All cDNA samples were diluted with nuclease-free water to the final cDNA concentration of 20 ng/μl.

Quantitative PCR was performed with The StepOne™ Real-Time PCR System or QuantStudio™ 3 System (Applied Biosystems™) using 5 μl diluted cDNA, 0.3 μM of the forward and reverse (5 μl) primer mix, and 12.5 μl 2× Thermo Scientific Maxima SYBR Green/ROX qPCR Master Mix (Thermo Scientific) and 2.5 μl of nuclease-free water in a reaction volume of 25 μl. The RT-qPCR reaction mix was denatured at 95 °C for 10 min and then subjected to 40 amplification cycles (15 s denaturation at 95 °C, 30 s annealing at 60 °C and 30 s extension at 72 °C) following manufacturer’s protocols for the SYBR Green Master Mix.

The reactions were run in triplicates. The specificity of the qPCR products was assessed by melting-curve analysis. No-template controls (NTC) did not record any positive Ct values. The Ct values of *CMAS* and *GAPDH* in surgically resected human brain tissue RNA samples were defined by TaqMAN qPCR assay. The following primers were used: *CMAS* (Hs00218814_m1, #4331182, Thermo Fisher Scientific) and *GAPDH* (Hs02758991_g1, #4331182, Thermo Fisher Scientific).

### Integrity value correction for difference in amplification efficiency

To estimate amplification efficiency, seven 5-fold dilutions were prepared using constructed linearized plasmids or prepared mouse or human cDNA and RNAse/DNAse free water. For plasmid dilution series, the highest concentration was 1.04×10^8^ plasmids per reaction. For cDNA dilution series the highest concentration was 170 ng of cDNA per 25 µl q-PCR reaction. RT-PCR was performed as described above. Acquired mean Ct values were plotted against concentration logarithm by 5 (the number of dilution factor) using Microsoft Excel Spreadsheet Software. Next, a linear regression curve was generated and the slope of the trend line was calculated. Amplification efficiency was estimated using the following equation:

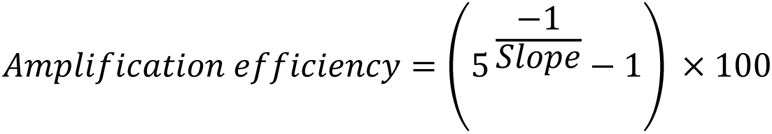

Then the amplification factor was calculated and corrections for the expression were made as described by Damgaard and Treebak^12^.

### Statistical analysis

GraphPad Prism 8 software was used for statistical analysis. R-squared test was applied to determine the strength of the relationship between the linear model and the dependent variables. Prism’s linear regression analysis was used to compare the slopes and the intercepts of two linear regression lines. Concordance between datasets was estimated through Pearson correlation. Outliers were defined as exceeding three interquartile ranges and removed from the analysis. Shapiro-Wilk test was used to assess the normality of the datasets. Paired *t*-test was used to compare RIN to 5’:3’ integrity values.

## Acknowledgements

This research was supported by the European Regional Development Fund under grant agreement No 01.2.2-CPVA-V-716-01-0001 with the Central Project Management Agency (CPVA). We would like to thank Dr. Daiva Dabkeviciene (Institute of Biosciences, Vilnius University, Lithuania) for helpful comments on statistical data analysis. We are grateful to Dr. Etienne Hezolg for sharing vGLUT1^mVenus^ mouse line. We thank Prof. S. Klimasauskas (Department of Biological DNA Modifications, Institute of Biotechnology, Vilnius University) and Dr. Ausra Sasnauskiene (Department of Biochemistry and Molecular Biology, Institute of Biosciences, Vilnius University) for sharing their equipment. Lastly, we thank patients who agreed to donate their post-surgical tissue for research.

## Author contributions

D.B. carried out major part of the experiments, performed data analysis and visualization, wrote the original draft of the article. K.M. and U.K. prepared and analysed the majority of human RNA samples. J.D. and E.B. provided the equipment and collaborated on FASS experiments. S.K. prepared plasmids. G.L. and S.R provided surgically resected human brain samples. U.N. supervised the whole study and the writing of the article. All authors approved the final version of the manuscript for submission. The manuscript has not been accepted or published elsewhere.

## Declaration of interests

The authors declare no competing interest.

## Notes

### Competing Interest Statement

The authors have declared no competing interest.

