## Supplementary figure S1, S2 and S3; Supplementary table S1 and S2 for "Integrity assay for messenger RNA in mouse and human brain samples and synaptosomal preparations"

Supplementary material

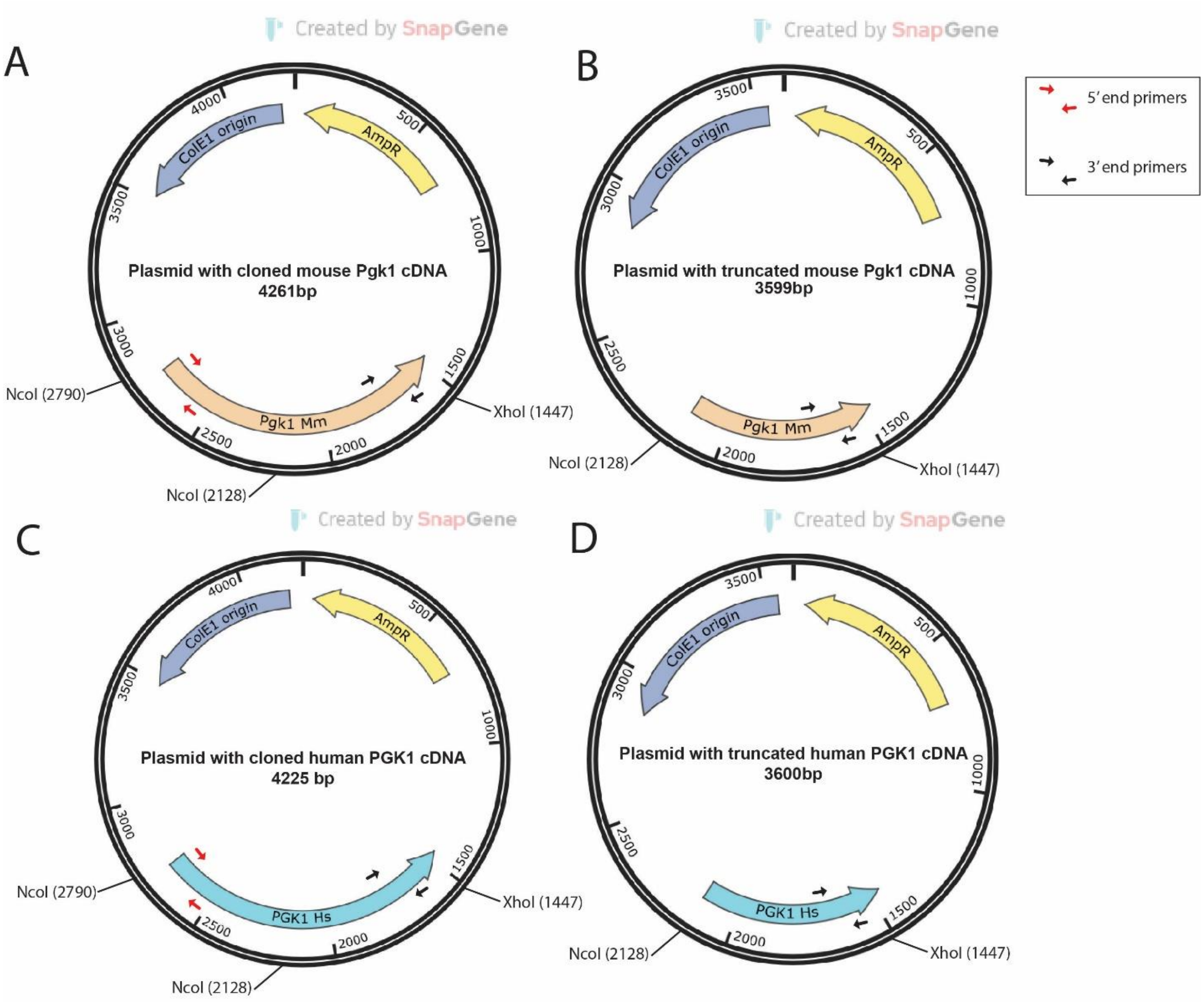


**Supplementary Figure S1 Plasmids with cloned *Pgk1* cDNA. a** Plasmid with cloned mouse *Pgk1* cDNA. **b** Plasmid with truncated mouse *Pgk1* cDNA. **c** Plasmid with cloned human *PGK1* cDNA. **d** Plasmid with truncated human *PGK1* cDNA. 5':3' assays primers are shown as red and black arrows.

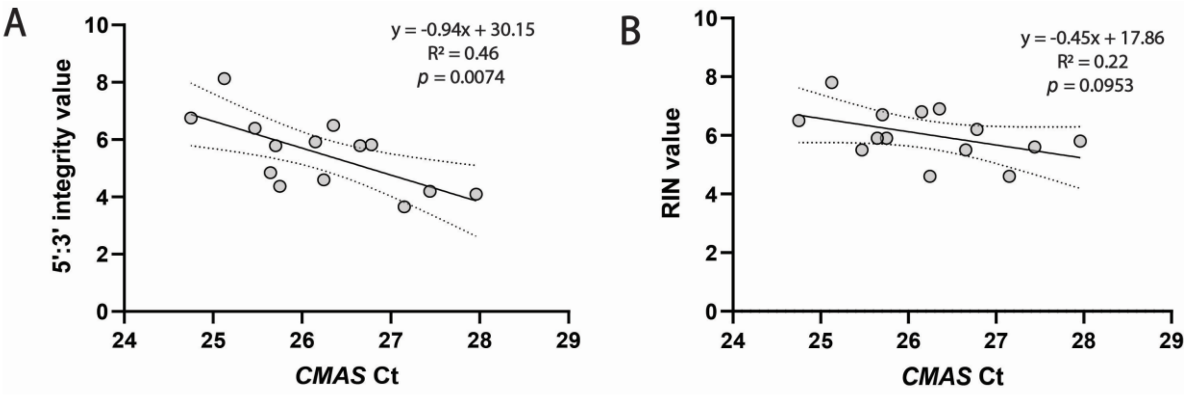


**Supplementary Figure S2 5':3'assay reflected transcript quality in RNA samples better then RIN value. A** There was a moderate linear correlation between the human brain tissue RNA sample 5':3' integrity value and Ct values from qPCR for *CMAS* expression while the correlation with RIN value was not significant (**B**)

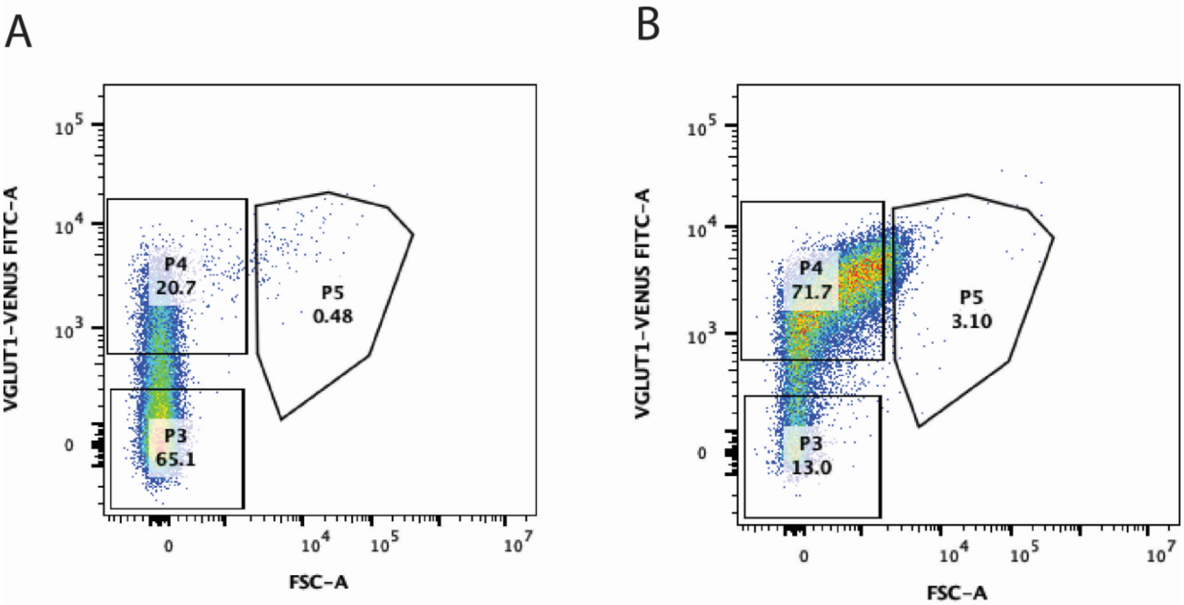


**Supplementary Figure S3 Analysis of synaptic particle distribution before and after sorting. A** In a crude synaptosome sample there was ~20% mVenus+ synaptosomes. B After FASS of mVenus+ excitatory synaptosomes, an enriched sample was prepared, where excitatory synaptosomes comprised more than 70 % of all particles.

**Supplementary table S1.** Donor tissue used in the study.

| ID | Sex | Age, years | Region of origin | Diagnosis |
|---|---|---|---|---|
| #02 | Female | 10 | Cortex | Hemisphere glioma |
| #03 | Male | 61 | Cortex | Supratentorial ependymoma |
| #04 | Male | 46 | Cortex | GBM |
| #05 | Female | 48 | Cortex | Epilepsy with hippocampal changes |
| #06 | Female | 44 | Cortex | Astrocytoma |
| #07 | Male | 40 | Cortex | GBM |
| #08 | Female | 28 | Cortex | Epilepsy with hippocampal sclerosis |
| #08a | Female | 28 | Hippocampus | Epilepsy with hippocampal sclerosis |
| #09 | Male | 23 | Cortex | Epilepsy |
| #09a | Male | 23 | Hippocampus | Epilepsy |
| #10a | Female | 29 | Hippocampus | Epilepsy |
| #11 | Male | 44 | Cortex | Oligodendroglioma |
| #12a | Female | 27 | Hippocampus | Epilepsy with hippocampal sclerosis |
| #12 | Female | 27 | Cortex | Epilepsy with hippocampal sclerosis |
| #13 | Male | 40 | Cortex | Epilepsy |
| #13a | Male | 40 | Hippocampus | Epilepsy |
| #14 | Male | 55 | Cortex | Oligodendroglioma |

**Supplementary table S2.** The comparison of 5':3' integrity values calculated without the primer amplification efficiency correction and the corrected 5':3' integrity values compared to RIN values. Original integrity values significantly differed from RIN ($p < 0.0001$, paired *t*-test), while corrected 5':3' integrity values and RIN were comparable ($p = 0.029$, paired *t*-test). The 5':3' integrity values and RIN were evaluated in RNA samples isolated from surgically resected human brain. List of samples and their clinical description is presented in Supplementary table S1.

| **Sample ID** | **5':3' integrity value** | **5':3' corrected integrity value** | **RIN** |
|---|---|---|---|
| #02 | 2.3 | 4.6 | 4.6 |
| #03 | 2.5 | 6.4 | 6.0 |
| #04 | 2.2 | 3.7 | 4.6 |
| #05 | 2.4 | 4.1 | 5.8 |
| #06 | 3.6 | 6.6 | 7.1 |
| #07 | 2.8 | 6.5 | 6.9 |
| #08 | 2.7 | 5.9 | 6.8 |
| #09a | 2.9 | 5.8 | 6.7 |
| #09 | 2.7 | 5.8 | 6.2 |
| #10a | 2.3 | 4.4 | 5.9 |
| #11 | 2.7 | 5.8 | 5.5 |
| #12a | 3.4 | 8.1 | 7.8 |
| #12 | 3.1 | 6.8 | 6.5 |
| #13 | 2.9 | 6.4 | 5.5 |
| #13a | 2.6 | 4.8 | 5.9 |
| #14 | 1.9 | 4.2 | 5.6 |
